# Efficient precision genome editing of *Chlamydomonas reinhardtii* with CRISPR/Cas

**DOI:** 10.1101/2022.07.24.501283

**Authors:** Adrian P. Nievergelt, Dennis R. Diener, Aliona Bogdanova, Gaia Pigino

## Abstract

CRISPR/Cas genome engineering in the unicellular green algal model *Chlamydomonas reinhardtii* has until now only been applied to targeted gene disruption, whereas scar-less knock-in transgenesis has generally been considered infeasible. We have developed highly efficient homology-directed knock-in mutagenesis in cell-walled strains of *Chlamydomonas*. Our method allows scarless integration of fusion tags and sequence modifications of near arbitrary proteins without need for a preceding mutant line.

## Main

The green unicellular alga *Chlamydomonas reinhardtii* is a popular model organism for topics ranging from structure and function of cilia and basal bodies^1,2^, chloroplast biogenesis^3^ and photosynthesis^4^ to circadian rhythm and share many protein homologues with higher eukaryotes^5^. *Chlamydomonas* offers a number of highly desirable traits for experimental work^6^: the cells grow readily by vegetative division in minimal media, both in suspension as well as on solid media, allowing for the generation of large biomass as well as simple isolation of clonal cells. Cell lines of opposite mating types can be crossed by sexual reproduction to combine genetic traits^7^. Additionally, vegetative cell lines are haploid, which facilitates genetic editing as only one successful alteration of a gene is necessary^8^. Finally, *Chlamydomonas* tends to integrate exogenous genetic material via random insertion into its genome, primarily by nonhomologous end joining (NHEJ). This property has allowed for the realisation of the Chlamydomonas Library Project (CLiP)^9^, the primary source of insertional mutants for the field. Traditionally, such mutants would be rescued by introduction of a recombinant copy of the originally disrupted gene. Such rescue experiments also allow for the introduction of fusion tags for biochemical or optical interrogation. However, rescue experiments become increasingly difficult for larger genes and will additionally disrupt a new, random, and thus unknown, part of the genome. Pioneering work by Greiner^10^, Shin^11^ and Picariello^12^ has established targeted insertion of exogenous DNA by the CRISPR/Cas9^13^ system for gene disruption in *Chlamydomonas*, but precision genetic editing by homology directed repair (HDR), a requirement for endogenous tagging, has so far been unattainable. Here, we demonstrate highly efficient and robust endogenous tagging of *Chlamydomonas* genes.

Based on the previous methodological developments^12^ we have established CRISPR/Cas9 based gene disruption in our work. To this end, we have constructed a set of vectors that allow for easy assembly of insertional cassettes by adding homology arms to an insert containing a selectable marker by Gibson assembly. We provide resistance markers to Paromomycin^14^, Nourseothricin^15^, Blasticidin S^16^ and Spectinomycin^17^ under the control of the RbcS2-promotor/RbcS2-1 Intron/RbcS2-terminator regulatory elements. The final insertional constructs are digested with type IIs restriction endonucleases and delivered together with Cas9 ribonucleoprotein (RNP) complexes by electroporation into heat shocked cells with autolysin-stripped cell walls (see Fig.1A).

**Figure 1:**
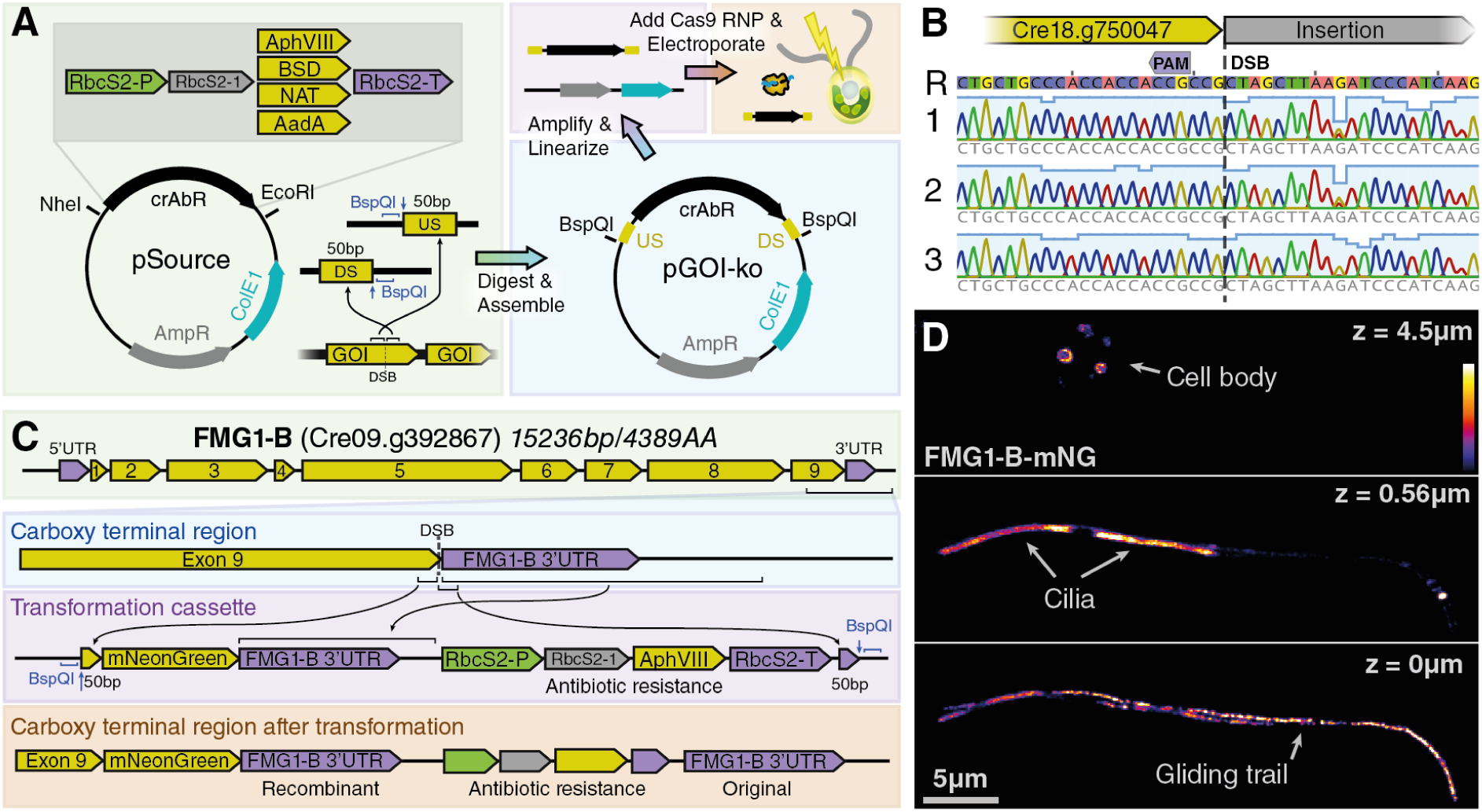
CRISPR/Cas9 enables precision genomic editing of *Chlamydomonas reinhardtii*. **A)** A selection of source vectors with commonly used *Chlamydomonas* antibiotic resistance markers (crAbR) allows for simple integration of synthetic homology arms corresponding to the flanks up-stream (US) and downstream (DS) of the designed Cas9 double strand break (DSB). **B)** Representative Sanger sequencing traces showing repeatable scar-less homology directed integration of a targeted cassette insertion. **C)** Construction of a cassette for a fluorescent mNeonGreen fusion to the carboxy terminus of FMG1-B. D) Spinning disk confocal sections of FMG1-B-mNeonGreen cell line, showing brightly labelled cilia (middle), trails of FMG1B-mNG due to gliding on the glass slide (bottom) and fluorescent intracellular vesicles (top).

In our quest to increase the insertional efficiency we have found that one of the most deciding factors is the purity of the DNA used during transformation. As such, the kit used for preparation of the DNA is crucial and we further reduce endotoxins in the purified DNA by an additional column purification step. The requirement for highly pure donor DNA is well established in other eukaryotic systems^18^. Additionally, we find that the transformation procedure is sensitive to a number of key parameters, most notably concentration of donor DNA and RNP, density of cells during heat shock treatment and cell wall removal, as well as the electroporation system used (see methods for optimal values). In our hands, the fully optimised process typically results in about 90% of the colonies expressing the selectable marker with inserts at the location of the intended double strand break (DSB) induced by Cas9 (see SFig1).

Importantly, the quantification of the lengths of the resulting insertions generally exhibit one discrete size that appears more frequently than others, indicating a favoured outcome of an insertion experiment. Indeed, Sanger sequencing of these insertions reveals almost all of these insertions to be the result of perfect homology directed repair (see Fig1B). We thus reasoned that this methodology, which has been primarily used for gene disruption in *Chlamydomonas*, can be directly extended to creating functional knock-in edits at endogenous loci. As proof of principle we chose a cut site close to the carboxy terminus of the flagellar major glycoprotein FMG1-B (Cre09.g392867v5)^19^. Using our resistance marker constructs, we extended the flank upstream of the cassette not only by a 50bp homology arm, but additionally by the short sequence starting at the DSB, up to (but not including) the stop codon, sequence coding for a fluorescent fusion tag (mNeonGreen) with linker followed by a full duplication of the 3’UTR of FMG1-B (see Fig1C). When electroporated into cells together with the corresponding Cas9 RNP and selected on antibiotics, this construct results in colonies of which about 20% consist of cells with bright green fluorescent flagella, as expected of a labelled flagellar coat protein (see Fig1D, Videos S1,S2). This result proves the feasibility of HDR driven endogenous knock-ins in *Chlamydomonas*.

Having established that knock-in fusions are feasible, we realised that the limiting factor for creating cell lines with endogenous tags is selecting the correct colonies, and set out to remove this bottle-neck by developing a highly robust screening procedure based on quantitative PCR (qPCR). A qPCR based approach is high-throughput compatible and drastically reduces the number of DNA gel electrophoresis steps: inserts can be directly detected based on amplification curves (see Fig2A, FigS2) and homology directed repair inserts can be identified by high-resolution melting analysis of the products (see Fig2B). The resulting candidates can then be verified by Sanger sequencing and phenotypic analysis. However, qPCR is sensitive to PCR inhibitors found in crude cell lysate and we have found multiple commercially available master mix reagents to fail to amplify even short amplicons in our hands. On the other hand it is crucial to be able to use crude lysate as a template for genotyping PCR, as purification would be prohibitively work intensive. We solved this problem by developing our own qPCR master mix based on an inhibitor tolerant PCR master mix together with a low inhibition qPCR dye (see methods). Using this custom master mix we are able to robustly amplify the flanks of insertional mutants over entire plates, while additionally being able to perform post-PCR high-resolution melting analysis of the products.

**Figure 2:**
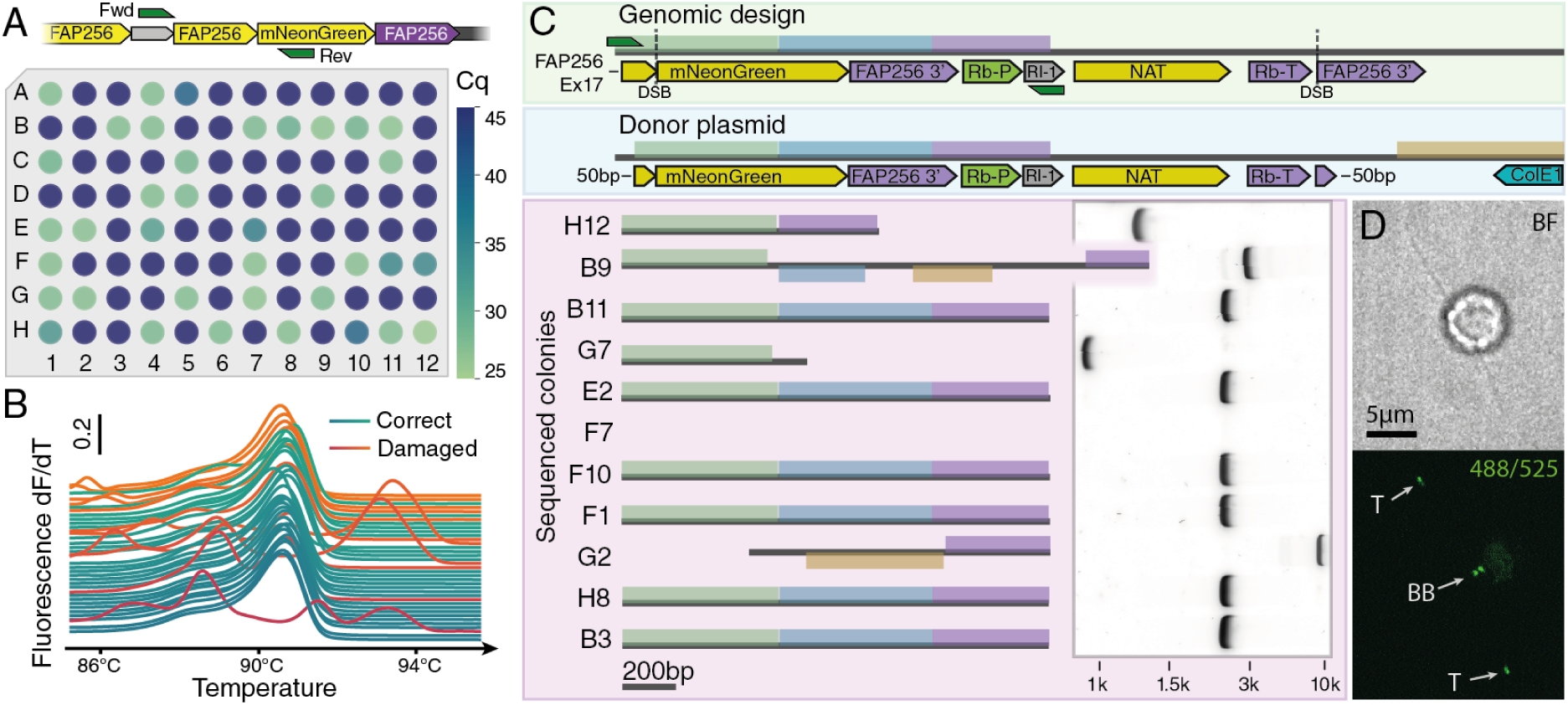
High-throughput screening allows rapid pre-selection of knock-in colonies with homology-directed repair insertions. **A)** Quantitation cycle (Cq) based screening for insertion by primers flanking the intended insertional junction. Colonies with low cycle numbers (green) are likely to have the insert. **B)** High-resolution melting (HRM) analysis of the products of A) allows for direct classification into wanted homology-directed repair insertions (green curves) and unwanted incorrect insertions (red curves). **C)** Simplified Mauve^20^ alignment of insertion sequences of 10 colonies selected based on HRM analysis in B) to the genomic design and the donor plasmid used to generate the strains. Large-scale rearrangements with the donor plasmid are seen in clones B9 and G2. Clones B11, E2, F10, F1, H8 and B3 are identical and exhibit a genotype as-designed. Amplicons with indicated primers (green) used for sequencing shown in agarose gel electrophoresis (See also figure S3). D) Bright-field (top) and fluorescence (bottom) micrographs of a ciliated FAP256-mNeonGreen knock-in cell confirm the expected localization at the ciliary tip (T) and the basal body (BB).

Strikingly, different colonies that have integrated a fluorescent fusion knock-in by HDR do not invariably exhibit the expected fluorescence signal. In addition to integrating the intended cassette, we find that these cases also exchange fragments of the non-selective part of the insert by recombination of micro-homology domains, thus rendering the insert non-functional (see Fig2C). In contrast, correct clones all exhibit the expected fluorescent signal (see Fig2D). As such, we recommend verifying the insertions of all new cell lines by Sanger sequencing.

Aside from defects during integration of the donor DNA, CRISPR/Cas9 is known to cause off-target defects^21^. Additionally, *Chlamydomonas* can integrate parts of the donor DNA in non-intended loci of the genome^9^. In order to search for these unintended changes and gauge the frequency at which this happens when constructing fusion knock-in lines we performed whole genome sequencing by short reads on four single knock-ins as well as one double knock-out and compared the resulting assemblies to the background genome. While we did not detect any off-target effects caused by Cas9, we have found random non-homologous integration events of vector fragments, as is typical of *Chlamydomonas* in one knock-in line as well as the double knock-out line. This result is consistent with the ∼10-20% of resistant colonies that do not have an insertion at the DSB and which must have inserted the resistance by random insertion. Thankfully, such unwanted insertions are often not problematic and can be easily removed by back-crossing.

In conclusion, we show that knock-in constructs for endogenous tagging are not only possible, but a feasible route to genetic engineering in *Chlamydomonas*. We believe that our approach has significant advantages to traditional rescue-type constructs, such as single-copy integration as well as the possibility of leaving edited genes under the control of their endogenous transcriptional elements. We expect this method will find wide-spread adaptation in the field to accelerate and enable work with this beautiful and versatile model organism.

## Methods

### Donor DNA construction and cloning

The nourseothricin resistance cassette was ordered as a gene fragment from Genscript. Blasticidin S resistance was ordered as a gene fragment from Eurofins. AphVIII and AadA genes were amplified from pE345 and pALM32 plasmids, respectively. All resistances were cloned into a high-copy vector containing the RbcS2 promoter and intron as well as the RbcS2 terminator using either restriction cloning or Gibson assembly.

All Gibson reactions were assembled with 2x NEB Hifi assembly master mix and incubated for 30min at 50°C. Fragments below 200bp were added in two-fold molar excess. Assembled DNA was transformed into chemically competent *E*.*coli* (GB06 or Dh5a). Mini-prep from LB liquid cultures were done using ZymoPure MiniPrep kit. Midi-prep for transformation into *Chlamydomonas* was done using ZymoPure MidiPrep kit including EndoZero columns for endotoxin removal. Donor DNA prepared by MidiPrep was digested with the corresponding restriction enzyme (usually NEB BspQI) according to manufacturer’s instructions and column purified by Zymo Clean and Concentrator 25 kit to ∼400ng/µL.

### Guide RNA selection and RNP preparation

crRNAs were designed in Geneious Prime and scored for off-targets and activity. Guides were selected to be close to the desired locus, have no detected off targets and ideally have an activity score above 0.5. gRNAs and tracrRNA were purchased from IDT. crRNA and tracrRNA were reconstituted to 100µM in nuclease-free duplex buffer. 10µL of crRNA and tracrRNA were mixed, heated to 95°C for 2 minutes and then removed from the heat block to cool to RT. RNPs were assembled by diluting 1.8µL AltR Cas9 v3 and 2µL of gRNA into 16µL of duplex buffer at RT.

### Transformation

Source cells were resuspended from a plate into 100µL TAP medium^8^ and then spread on 100mm plates with TAP in 1.5% Agar and grown in a 14h light/10h dark cycle for 3-4 days until a lawn formed. Resulting cells were resuspended in 1mL TAP in a 1.5mL tube and pelleted. All centrifugations were 600RCF for 2 minutes. The supernatant was carefully aspirated, and the pellet resuspended in 100µL. This suspension was aliquoted into fresh 1.5mL tubes to result in cell pellets of 10µL. Each tube was treated 3 times with gamete autolysin to remove cell walls, where the third wash was combined with 30 minute heat shock at 40°C^12^. Cells were then washed 3 times in 1.4mL TAPS (TAP+40mM sucrose) and finally resuspended in 80µL TAPS and realiquoted at 40µL per tube into fresh 1.5mL tubes. To each tube 4µL of RNP solution and 1.2µg of linearized donor DNA was added. Cells were electroporated in 4 reactions of 10µL with the Neon electroporation system at 3200V for 12ms and 3 pulses. Finished reactions were pipetted directly into 1mL TAPS in 24-well plates and left for recovery overnight. Finally, cells were concentrated by centrifugation and spread onto TAP agar plates with the corresponding antibiotic. Plates were incubated in constant light until colonies formed, between 3-7 days.

### Screening qPCR and sequencing

Colonies from antibiotic plates were picked into 96 well plates and kept in constant light until the wells turned green. 10µL of each well were subsequently mixed with 10µL of Lucigen QuickExtract buffer and heated to 65°C for 6 minutes, then 95°C for 2 minutes. qPCR plates were filled with reactions consisting of 3µL Invitrogen Platinum 2 HS mastermix, 0.15µL of 10µM primer mix targeting the desired insert, 0.5µL template from the previous step, 0.3µL EvaGreen Plus and 2.05µL of 2M Betaine. Plates were cycled in a Roche Lightcycler 96 according to the manufacturer’s recommendations. Curves were analysed for quantification cycle Cq and high resolution melting curves by LightCycler software.

The insert of promising wells was amplified by PCR, using Invitrogen SuperFi2 Master Mix with final 1M Betaine in the reaction. PCR products were sized by 1.2% agarose gel electrophoresis in 10mM lithium-acetate-borate buffer, stained with Biotinum GelGreen and visualised on a Typhoon FLA 9500 imager. Finally, PCR products were column purified and sent for Sanger sequencing.

### Whole genome sequencing

50mL of dense liquid culture was pelleted by centrifugation at 1200RCF for 10 minutes, followed by 4 steps of extraction with phenol-chloroform-isoamyl alcohol (25:24:1). DNA was precipitated by addition of 2 volumes of cold ethanol, pelleted at 16kRCF for 5 minutes, then air dried and resuspended in 200µL TE overnight.

Adapters were trimmed with trim_galore and paired reads were mapped to the CC1690 assembly from NCBI using bowtie2. Variants were called using deepvariant and subsequently filtered to the ±10kb vicinity of potential off-target sites identified by blast using a high e-value cut-off. Variants were further filtered using the Ensembl Variant Effect Predictor to variants overlapping gene sequences. Variants that were also detected in the background strain were removed.

### Optical microscopy

Spinning disk micrographs have been acquired with an Andor IX 83 equipped with a Yokogawa CSU-W1 spinning disk at a 50µm pinhole size through an Olympus 150x/1.45 U Apo oil immersion objective on an Andor iXon Ultra 888 resulting in a pixel size of 79nm. Z-stacks were acquired using a Prior NanoScanZ piezo stage in 281nm optical sections and subsequently deconvoluted with the spinning disk module of Huygens deconvolution. False colour mapping was applied with Fiji^22^.

## Acknowledgments

The authors would like to acknowledge M. Sarov, I. Reichardt-Gomez and J. Koellner from the genome engineering facility at MPI-CBG as well as T. Brown and the DRESDEN-*concept* Genome Center (DcGC), the sequencing facility and the FACS facility at MPI-CBG. We would also like to thank C. Peano and N. Alfonso from the Genomics Facility of Human Technopole. We would like to acknowledge support from the MPI-CBG light microscopy facility. A.P.N was supported by an EMBO Long--term fellowship under ALTF number 891-2018 as well as by an HFSP Cross-disciplinary fellowship with reference number LT000515/2019. We would like to acknowledge the European Research Council (ERC) under the European Union’s Horizon 2020 research and innovation program (grant agreement No. 819826) and the DFG grant GACR-DFG Cooperation 2019 No.PI1218/3-1 to G.P.

## Author contributions

Conceptualization, A.P.N, G.P; Methodology, A.P.N, D.D; Investigation, A.P.N, D.D, A.B; Data Curation, A.P.N; Visualisation, A.P.N; Formal Analysis, A.P.N; Writing – Original Draft, A.P.N; Writing – Review & Editing, A.P.N, D.D and G.P; Funding Acquisition, A.P.N and G.P; Resources, G.P; Supervision, G.P

## Data Availability

Plasmids and *C. reinhardtii* lines generated in this study will be made available to the Chlamydomonas Resource Center at the time of publication.

## Supplementary Information

**SFigure 1:**
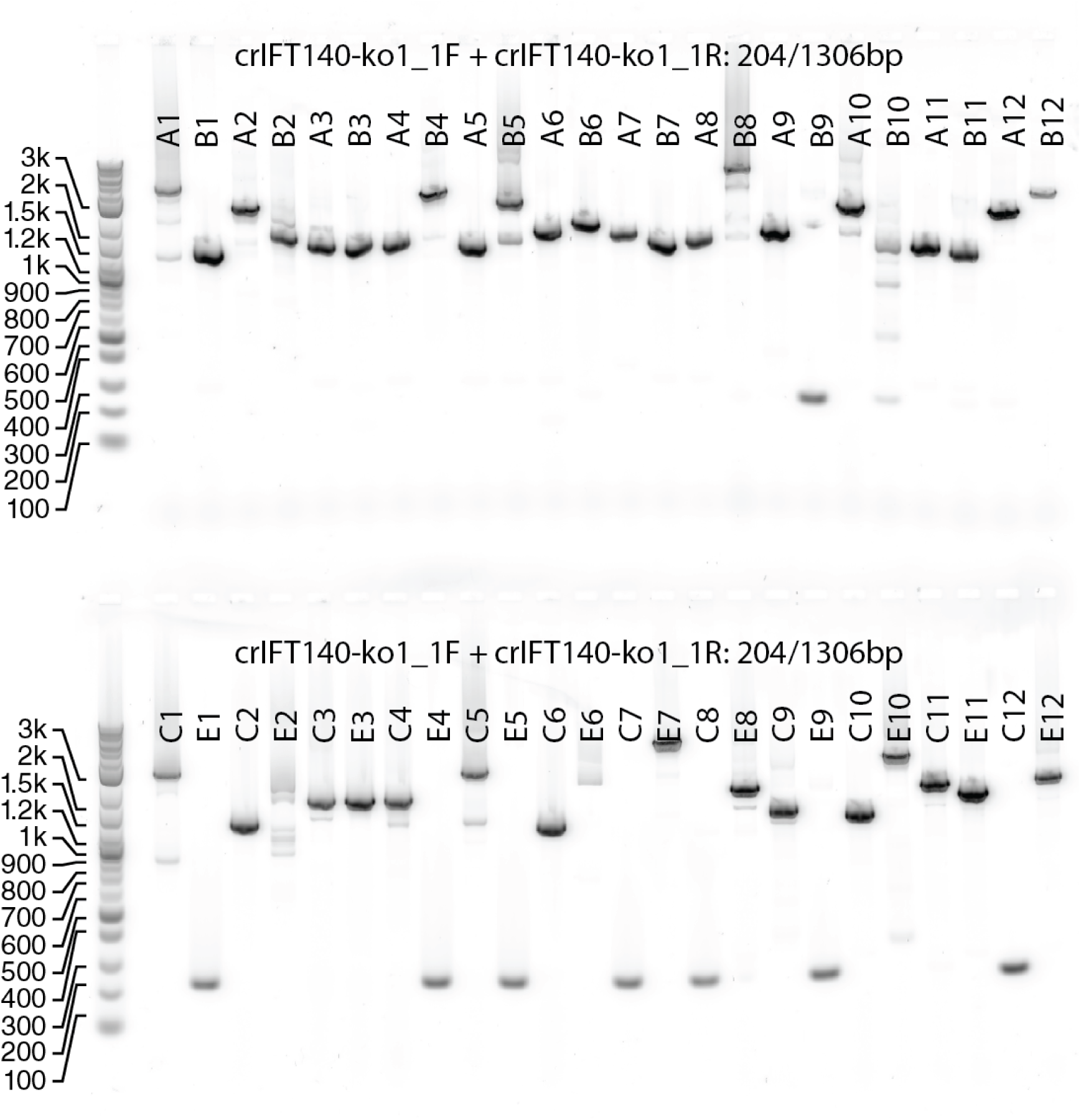
Annotated agarose gel electrophoresis of PCR products showing band upshifts due to insertion of a knock-out cassette in the IFT140 gene (Cre08.g362650). 8/96 bands show a wild-type band at 204bp, while the rest exhibit a variety of insertion sizes with a median around 1306bp expected for correct homology directed insertion.

**SFigure 2:**
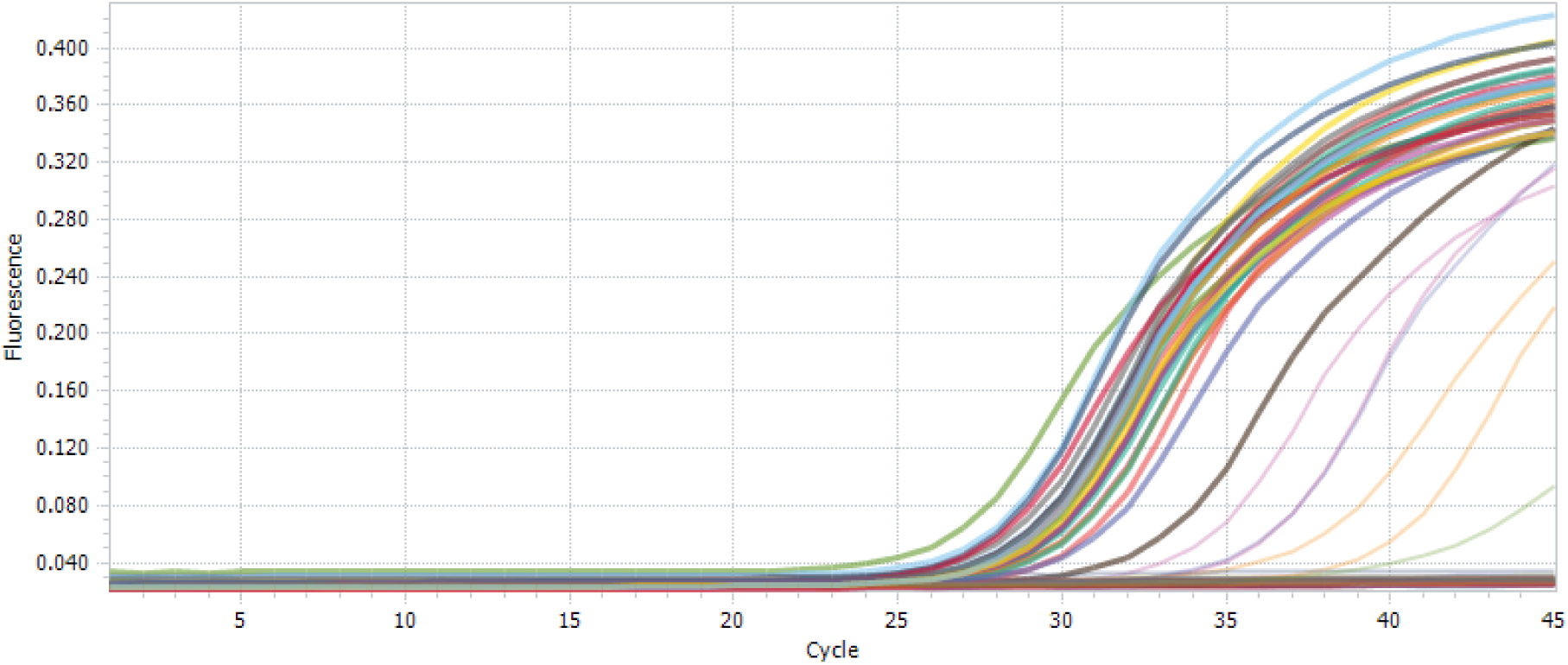
Raw qPCR amplification curves for plate map in figure 2A.

**SFigure 3:**
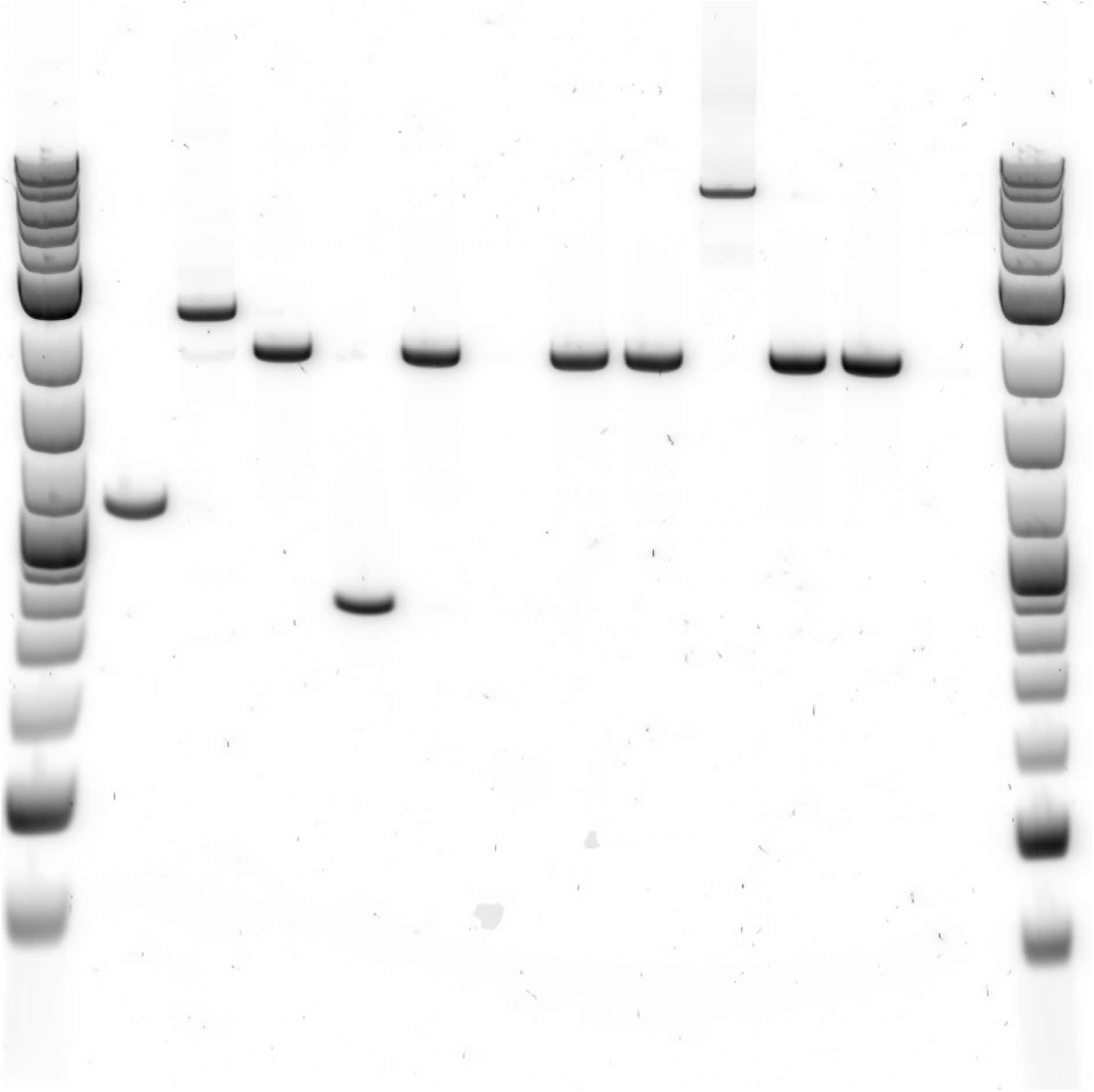
Original image for agarose gel electrophoresis shown in Figure 2c. Lanes are: 1,14 NEB 1kbp plus; 2-13: Genomic insert PCR products for colonies H12, B9, B11, G7, E2, F7, F10, F1, G2, H8, B3.

**SVideo 1:** Low-magnification fluorescence widefield time-lapse of FMG1-B-mNeonGreen fusion knock-in cells shows intracellular as well as ciliary fluorescence in swimming cells.

**SVideo 2:** High-magnification widefield time-lapse of FMG1-B-mNeonGreen fusion knock-in cells showing fluorescent cilia of cells either gliding on or swimming close to the glass surface.

